# BiRNA-BERT allows efficient RNA language modeling with adaptive tokenization

**DOI:** 10.1101/2024.07.02.601703

**Authors:** Md Toki Tahmid, Haz Sameen Shahgir, Sazan Mahbub, Yue Dong, Md. Shamsuzzoha Bayzid

## Abstract

Recent advancements in Transformer-based models have spurred interest in their use for biological sequence analysis. However, adapting models like BERT is challenging due to sequence length, often requiring truncation for proteomics and genomics tasks. Additionally, advanced tokenization and relative positional encoding techniques for long contexts in NLP are often not directly transferable to DNA/RNA sequences, which require nucleotide or character-level encodings for tasks such as 3D torsion angle prediction. To tackle these challenges, we propose an adaptive dual tokenization scheme for bioinformatics that utilizes both nucleotide-level (NUC) and efficient BPE tokenizations. Building on the dual tokenization, we introduce BiRNA-BERT, a 117M parameter Transformer encoder pretrained with our proposed tokenization on 28 billion nucleotides across 36 million coding and non-coding RNA sequences. The learned representation by BiRNA-BERT generalizes across a range of applications and achieves state-of-the-art results in long-sequence downstream tasks and achieves a performance comparable to 6× larger models in short-sequence tasks with 27×less pre-training compute. BiRNA-BERT can dynamically adjust its tokenization strategy based on sequence lengths, utilizing NUC for shorter sequences and switching to BPE for longer ones, thereby offering, for the first time, the capability to efficiently handle arbitrarily long DNA/RNA sequences. ^1^

## 1 Introduction

The introduction of encoder-only transformer models has revolutionized Natural Language Processing (NLP) by significantly improving our ability to extract deep semantic representations from text which can be used for a wide range of downstream tasks [9, 19]. This success has inspired researchers to apply the pretraining-finetuning paradigm to a wide range of topics beyond NLP, including biological sequence modeling with remarkable success [3,10,15,21,43]. However, transferring improvements from natural language processing (NLP) to biological sequences is not always straightforward. A key challenge arises from the need to model extremely long sequences for certain tasks, as biological sequences can range from tens to millions of nucleotides or amino acids. The standard transformer encoder architecture (e.g., BERT [9]) uses positional embeddings that limit the maximum input length to usually 512 or 1024, requiring truncation or inefficient workarounds for biological sequences.

In addition, despite recent advancements in methodological and architectural research efforts to scale up the length for BERT [4,12,22] by addressing the limited context window [23,33] or with more efficient tokenization methods [11,28], they have not yet been successfully adopted by bioinformatics without sacrificing performance or sample efficiency. The limited number of approaches to adopting new positional encoding and improved pretraining developed in NLP often fall short in biological sequencing. For example, Nucleotide Transformers (NT) [7] replace nucleotide-level tokenization (NUC) with non-overlapping *k*-mers, where *k*-mers represent subtokens of fixed length *k*, enabling the model to handle sequences that are *k* times longer while sacrificing granularity. This limitation is acknowledged in DNABERT-2 [43], where they demonstrate the poor sample efficiency of *k*-mer tokenization and propose adapting byte pair encoding (BPE) tokenization for DNA sequences. However, their approaches still suffer from granularity loss by combining common subsequences into one token, and the original paper does not provide a controlled comparison between NUC and BPE. Moreover, in domains like RNA or protein sequences where secondary structure prediction or granular alignment tasks are important, simply using BPE tokenization would prevent the trained foundation model from generating nucleotide- or amino acid-level predictions, which are essential for these additional tasks.

Following the success of *foundation models* – models pre-trained on vast amounts of unlabelled data – in protein sequences, building similar pre-trained models for DNA/RNA sequences has gained a significant attention from bioinformatics researchers. Notable foundation models for DNA include DNABERT [15], DNABERT-2 [43], and Nucleotide Transformer [7], all of which are BERT-style encoder-only models largely following the pretrain-then-finetune paradigm. Considerable progress has been made for RNA sequences as well. [3] proposed RNA-FM, a 100 million parameters model pre-trained on 23.7 million unannotated ncRNAs from the RNAcentral database [5] using the standard masked language modeling task and nucleotide-level tokenization (NUC). Uni-RNA [38] was pre-trained on a dataset of 1 billion RNA sequences for a spectrum of structural and functional downstream tasks. [42] proposed a Multiple sequence alignment (MSA)-based RNA language model (RNA-MSM), a BERT-style language model that utilizes a set of homologous sequences per forward propagation and produced attention maps and embeddings that have direct correlations to RNA secondary structure and solvent accessibility without supervised training. For downstream tasks, RNA-MSM finds homologous sequences of the input sequence from a stored database, computes the MSA matrix using RNAcmap [41], and uses information from the MSA as input to the model. However, obtaining the MSAs is a time-consuming procedure, for example, it takes RNAcmap on average 9 hours to obtain an MSA for one RNA sequence of length 60. [21] introduced RiNALMo, a 650M parameter model that incorporates algorithm and architecture improvements in NLP models, namely Rotary Position Encoding (RoPE) [33], FlashAttention2 [8], and SwiGLU [30], and is pre-trained on 36 million ncRNA sequences from RNAcentral [5], NT, Rfam [13], and Ensembl databases [6]. RiNALMo achieves state-of-the-art performance on a variety of downstream tasks such as splice-site and mean ribosome loading and exhibits stronger generalization than previous thermodynamics-based and deep learning models on secondary structure prediction.

All the aforementioned biological foundation models either use character-level (nucleotide-level) tokenization or BPE tokenization, each with its trade-offs. Approaches using nucleotide-level tokenization inevitably face long tokenized sequences, while approaches using BPE, such as DNABERT-2, do not cater to tasks requiring nucleotide-level granularity. Given these limitations, we propose **Bi**-tokenization **RNA BERT** (BiRNA-BERT) – a Transformer encoder model for RNA sequences pretrained on both NUC and BPE tokens of the same RNA sequences simultaneously. BiRNA-BERT uses Attention with Linear Biases (ALiBi) [23] which allows the context window to be extended without retraining and can dynamically choose between NUC and BPE tokenization based on the input sequence length. For shorter sequences, it utilizes NUC to capture fine-grained patterns, and for longer sequences, it switches to more efficient BPE tokenization to reduce memory requirements without truncating the input. This dynamic tokenization and context length expansion allows BiRNA-BERT to enable downstream tasks with arbitrarily long sequences and set state-of-the-art results on 4 out of 6 tasks of miRNA-lncRNA interaction dataset [39]. Along with sequence-level tasks, the dual tokenization approach allows us to conduct nucleotide-level analyses, such as RNA 3D torsion angle prediction [32], as additional concurrent downstream tasks.

We conduct further downstream ablation experiments on both RNA and DNA data to demonstrate that NUC tokens yield better downstream task performance than BPE when enough resources are available to handle the short sequences. In addition, the joint-pretraining strategy does not compromise nucleotide-level performance compared to models trained on only NUC tokens.

In summary, BiRNA-BERT demonstrates the effectiveness and importance of utilizing proper tokenization tailored to the unique demands of bioinformatics tasks. Pre-training models leveraging our proposed tokenization can achieve significant performance gains across a range of biological sequence analysis tasks. Our main contributions can therefore be summarized as follows:

1. We, for the first time, present dual tokenization pretraining – an effective approach that extends the effective context window of biological foundation models with efficient tokenization while retaining the ability to generate character-level embeddings.
2. Using dual tokenization and ALiBi, we train BiRNA-BERT which achieves absolute state-of-the-art results on long-sequence tasks and is comparable to 6× larger models on short-sequence and nucleotide-level tasks while being trained with 27× less pretraining compute. BiRNA-BERT can dynamically adjust the tokenization algorithm based on the sequence length and available compute.
3. We, for the first time, provide a rigorous mathematical analysis from information theoretic perspective about the absolute information loss of token compression with byte pair encoding. We show that NUC outperforms BPE on tasks where sequences are short enough to fit into GPU memory and the trade-off between sequence truncation and information compression works in favour of the later when long sequences are in consideration.
4. We validate the adaptability of our proposed scheme to other biological language models beyond RNA sequences by extending the idea of dual tokenization with AliBi encoding to DNA sequences too with BiDNA-BERT which shows very competitive performance to the DNA foundation models, even being 66x less compute heavy.

## 2 Methods

The schematic diagram of BiRNA-BERT is shown in Figure 1, highlighting the major challenges in RNA language modeling and demonstrating how BiRNA-BERT addresses these challenges. We describe the key components and training strategies of BiRNA-BERT in the subsequent sections. We first describe the motivation behind using relative positional encoding (ALiBi) for extrapolating the trained model to longer sequences in downstream tasks (Section 2.1). Then we discuss about the tokenization strategies for biological foundation models and applicability of using BPE tokenization for BiRNA-BERT (Section 2.2). In Section 2.3, we discuss the dual tokenization pretraining approach and its motivation. Finally, we describe the pretraining configurations and datasets used in Sections 2.4 and 2.5.

**Figure 1:**
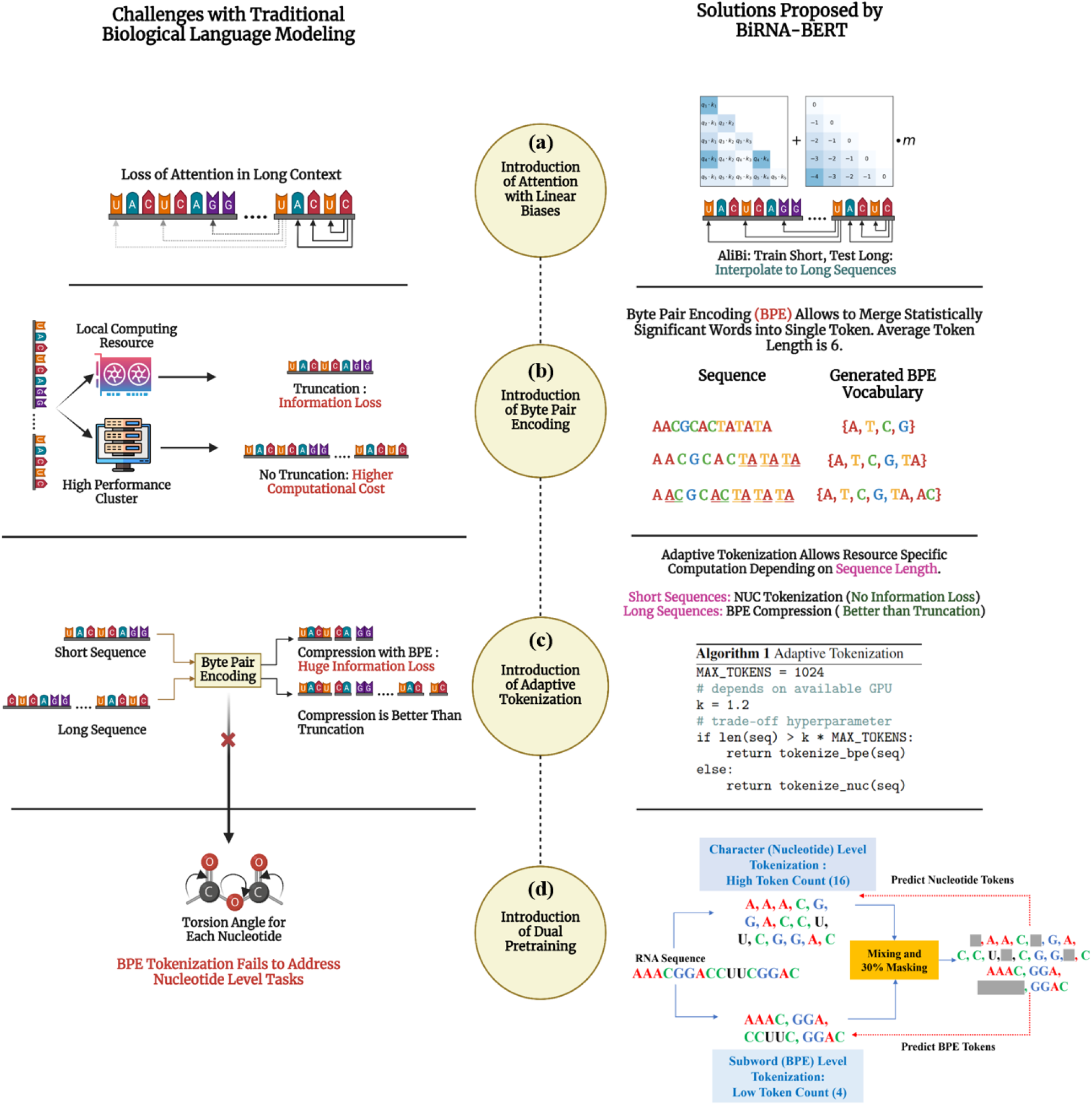
Schematic diagram of the overall architecture of BiRNA-BERT, illustrating its components designed to address key challenges in RNA language modeling. **(a)** BiRNA-BERT resolves the issue of loss of attention in long sequences by incorporating AliBi positional encoding. **(b)** For low-compute environments where large sequences cannot fit into memory, BiRNA-BERT employs BPE tokenization, which merges statistically significant residues into single tokens, allowing for longer context lengths within limited resources. **(c)** Although BPE compression is more effective than sequence truncation for long sequences, it leads to information loss and poor performance in shorter sequences. To address this, BiRNA-BERT uses adaptive tokenization: NUC tokenization is preferred for short sequences, while BPE tokenization is applied to long sequences, avoiding truncation. **(d)** To enable residue-level downstream tasks, BiRNA-BERT introduces dual pretraining, supporting both sequence-level and residue-level tasks simultaneously.

### 2.1 Positional Encoding in the Transformer Architecture

Since the attention mechanism is permutation-invariant, positional information must be explicitly added. There are two main strategies for encoding positional information - fixed and relative. In fixed positional encoding schemes such as sinusoidal [36] or learned embeddings [9], the positional information is a vector function of the position index within some predefined context length. Since the positional information is explicit in the form of a vector, these methods cannot extrapolate to context lengths beyond those seen in pretraining. Previous biological foundation models [3, 15] have mostly used fixed positional embeddings following the original BERT implementation. As a result, long sequences had to be truncated (usually to 1024 tokens) on downstream tasks. To overcome this challenge, we leverage relative positional encoding as shown in Figure 1(a). In relative positional encoding, the positional information is a function of the distance between two tokens. Popular algorithms for relative positional embeddings are T5 Bias [24], Rotary Positional Embedding (RoPE) [34], and Attention with Linear Biases (ALiBi) [23]. Recently, RiNALMo [21] utilized Rotary Positional Embedding (RoPE) but still truncated sequences to 1022 nucleotides since extending the context window using RoPE requires additional training to preserve performance. DNABERT-2 [43] was the first biological foundation model to use ALiBi, which when combined with BPE, allowed them to process sequences of up to 10000 nucleotides, significantly outperforming recurrent models such as HyenaDNA [20] on long-sequence tasks.

#### Attention with Linear Biases (ALiBi)

In contrast to complex methods such as T5 Bias and RoPE which are hard for models to extrapolate without continued pretraining, ALiBi simply reduces the attention score between two tokens by a scaler function of their distance. Let **Q** ∈ ℝ^*L*×*d*^, **K** ∈ ℝ^*L*×*d*^, and **V** ∈ ℝ^*L*×*d*^ represent the query, key, and value matrices, where *L* is the sequence length and *d* is the hidden dimension. ALiBi modifies the attention scores by adding a bias term, a function of the positional distance between the query and key elements. Specifically, for positions *i* and *j*, the linear bias term *b*_*ij*_ is defined as *b*_*ij*_ = *α·* |*i*− *j*|,where *α* is a hyperparameter. This results in modified attention scores 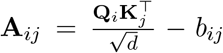 Since ALiBi explicitly penalized the attention scores between distance tokens, the *α* values must be carefully chosen to preserve long-context capabilities. The authors of ALiBi use different *α* values for different attention heads, uniformly sampled from the geometric progression 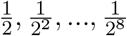. Intuitively, some attention heads specialize in aggregating local information while others preserve long-context information.

### 2.2 Tokenization Strategies for Biological Foundational Models

Nucleotide-level tokenization (NUC) has long been the dominant form of tokenization in biological sequence modeling [3,21], partially due to the need for NUC embeddings in secondary tasks for RNA and protein. An alternative form of tokenization is *k*-mer tokenization, where segments of *k* nucleotides are considered as one token. The motivation behind *k*-mer tokenization is to effectively capture the biological significance of RNA sequences, as certain *k*-mers, such as 3-mers (codons), directly correspond to amino acids, which are the building blocks of proteins. DNABERT [15] uses overlapping *k*-mers while models such as Nucleotide Transformers [7] use non-overlapping k-mers. However, these full sequence tokenization schemes are constarined by the memory limitation. Even with relative positional encoding, it is challenging to fit longer context length sequences into available memory. To tackle this challenge, we have employed Byte Pair Encoding (BPE) tokenization as shown in Figure 1(b). BPE [11, 28] is a subword tokenization technique that iteratively merges the most frequent pairs of bytes or characters to create new tokens, thereby reducing the vocabulary size to a fixed number.

Recently, DNABERT-2 [43] demonstrated the superiority of BPE tokenization over NUC and *k*-mer tokenization for DNA sequences. Since DNA lacks notable downstream tasks that necessitate NUC embeddings, BPE tokenization proved to be the most suitable approach for this context.

#### BPE Tokenization in RNA Sequence Modeling

In the context of RNA sequences, the variability and complexity of the nucleotide sequences pose a challenge for traditional tokenization methods. Fixed k-mer-based tokenization can result in an excessively large and sparse vocabulary of size 4^*k*^ + 5 [40], as it captures all possible k-mers regardless of their biological significance or frequency. BPE tokenization, on the other hand, leverages the statistical frequency of sub-sequences to create a more compact and meaningful vocabulary. By iteratively merging the most frequent pairs of sub-sequences, BPE ensures that commonly occurring patterns are represented by single tokens, while less frequent patterns are broken down into smaller units. This process is particularly beneficial for RNA sequences, where certain motifs and regions (e.g., hairpin loops, binding sites) occur frequently and are biologically significant. DNABERT-2 [43] tested the effect of the BPE vocabulary size on downstream tasks and empirically showed that a vocabulary size of 4096 is optimal. Following DNABERT-2, we use a vocabulary size of 4096 for the BPE tokenization.

We now present an example to illustrate the process of Byte-Pair Encoding (BPE) tokenization for RNA sequences. Consider an RNA sequence *S* = AUGGCUACUGCAUGCUAGUCA. Initially, the vocabulary consists of individual nucleotides: A, U, G, C. The BPE algorithm proceeds by iteratively merging the most frequent adjacent pairs of characters, as follows:

1. Identify frequent pairs: The algorithm identifies the most frequent pair of characters in the sequence. Suppose the pair “AU” appears most frequently.
2. Merge the pair: The pair “AU” is merged into a single token, updating the sequence:

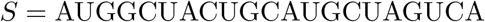
3. Repeat the process: The algorithm then identifies the next most frequent pair, say “GC”. Merging “GC” into a single token updates the sequence to:

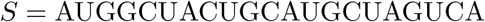
4. Continue merging: This process is repeated until the predefined vocabulary size is reached. At the end of the tokenization, the final sequence might be tokenized as:

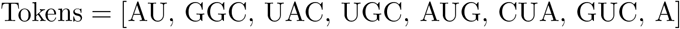

This approach effectively captures the frequent patterns within the RNA sequence, balancing the trade-off between compression and biological relevance.

### 2.3 Adaptive Tokenization and Dual Pretraining

BPE tokenization performs well for longer sequences, offering a significant advantage over fixed-length truncation, as discussed in the results section. However, for shorter sequences, this compression method can result in information loss (Section 3.5). To address this, we implemented an adaptive tokenization strategy: BPE tokenization is employed for long sequences, while NUC (nucleotide-level) tokenization is used for shorter ones, as illustrated in Figure 1(c). Another limitation of BPE tokenization is its inability to handle nucleotide-level tasks, such as torsion angle and secondary structure prediction, as BPE tokenization compresses multiple nucleotides into a single token. To overcome this, we propose a dual pretraining approach as shown in Figure 1(d), which allows a single model to learn both BPE tokens and NUC tokens effectively. The primary motivation behind training a single model with dual tokenization is memory efficiency. Although having separate BPE and NUC models is viable, we argue that a single model training on both is memory-efficient since only one set of weights needs to be on memory and can process longer sequences and larger batch sizes. Memory efficiency is crucial since the transformer architecture has *O*(*n*^2^) memory requirement which especially penalizes long sequence tasks. In addition to BiRNA-BERT, We train two more models BPE-Only-BERT and NUC-Only-BERT to show that dual tokenization does not affect downstream performance. Performing BPE tokenization on long sequences effectively compresses the sequence with average BPE token length 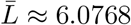. We show in Section 3.5 that the empirical per-character entropy ratio is:

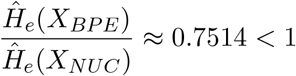

where Ĥ_*e*_(*X*_*BPE*_) is the average character-level entropy of the BPE representation of a sequence and Ĥ_*e*_(*X*_*NUC*_) is the per-character entropy for nucleotide tokenization. Therefore, BPE tokenization is essentially a trade-off between information compression and computational efficiency,

Given an RNA sequence **r** = (*r*_1_, *r*_2_, …, *r*_*n*_), we apply two types of tokenization: BPE and NUC.

- **BPE Tokenization:** The BPE tokenized sequence is **r**_*BPE*_ = (*β*_1_, *β*_2_, …, *β*_*m*_) with a vocabulary size of 4096.
- **Nucleotide-Level Tokenization:** The nucleotide tokenized sequence is **r**_*Nuc*_ = (*v*_1_, *v*_2_, …, *v*_*n*_) with a vocabulary size of 4.

We randomly mask tokens in both **r**_*BPE*_ and **r**_*Nuc*_ for Masked Language Modeling (MLM) training:

- **Masked BPE Sequence:** 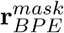 where a subset of *β*_*i*_ is replaced with a mask token [*MASK*].
- **Masked Nucleotide Sequence:** 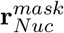 where a subset of *v*_*i*_ is replaced with a mask token [*MASK*].

The total loss *L*_*total*_ is the sum of the BPE and nucleotide losses:

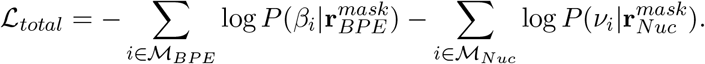

The parameters of the BERT model, Θ, are optimized to minimize the total loss:

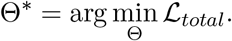

### 2.4 Pretraining Configuration

We pretrain four BERT-based RNA models to investigate the effect of dual tokenization: BPE-Only, NUC-Only, BiRNA-BPE, and BiRNA-NUC

We train our language model under three configurations:

1. **BPE-Only:** In this configuration, we tokenize the pre-training sequences only with Byte Pair Encoding (BPE). Since BPE tokenization unifies the most common subsequences into a single token, this configuration resulted in the least number of training tokens: 4.343 billion tokens. Training this version required 3 hours on 8× 3090 GPUs.
2. **NUC-Only:** In this version, we tokenize each sequence at the individual nucleotide level. This method resulted in 27.87 billion training tokens, and training took 45.1 hours on 8× 3090 GPUs.
3. **BiRNA:** In this configuration, we combine both BPE and NUC tokenizations to pretrain the RNA language model. This approach resulted in 32.254 billion tokens and required 48.42 hours of training on 8× 3090 GPUs. During inference, we utilize two methods:

- **BiRNA-BPE:** Inference is performed using BPE tokenization.
- **BiRNA-NUC:** Inference is performed using NUC tokenization.

We use the MosaicML framework [22] for efficient training. We pretrain all models with a learning rate (LR) of 2 × 10^−4^, a warmup ratio of 0.06, a linear rate scheduler that decreases to 0.02 of the starting LR, and a batch size of 200 per device, totaling 1600 samples per batch across eight Nvidia RTX 3090 GPUs. Due to the investigative nature of our work and compute constraints, we pretrain each model for only 1 epoch. The training configurations are summarized in Table 1.

**Table 1:** Pretraining details for various models.

| Model | Train Tokens | Train Time | Hardware |
| --- | --- | --- | --- |
| BPE only | 4.384B | 3 hours | $8 \times 3090$ |
| NUC only | 27.87B | 45.1 hours | $8 \times 3090$ |
| BiRNA-NUC | 32.254B | 48.42 hours | $8 \times 3090$ |

### 2.5 Pretraining Dataset

For the pretraining phase, we selected datasets comprising both ncRNA and mRNA sequences due to their complementary roles in cellular processes. ncRNA sequences, despite not encoding proteins, are integral to regulating gene expression and maintaining cellular functions. Conversely, mRNA sequences act as templates for protein synthesis, having a direct impact on the cell’s proteome.

#### Non-coding RNA Sequences

To assemble a robust dataset of ncRNA sequences, we utilized the RNAcentral database [5]. RNAcentral serves as an exhaustive repository, aggregating data from multiple ncRNA databases, thereby providing a wide array of ncRNA sequences. The dataset from RNAcentral encompasses approximately 36 million RNA sequences, collectively containing 26.42 billion nucleotides. This extensive compilation allows for a thorough representation of the ncRNA landscape, facilitating effective model training.

#### Coding RNA Sequences

For coding RNA sequences, we turned to the RefSeq database ^2^. RefSeq is a well-curated collection of sequences, providing high-quality mRNA sequences across various species. For each species cataloged in RefSeq, we extracted the mRNA sequences, culminating in a dataset of 532,852 mRNA sequences. These sequences have an average length of 4,175 nucleotides, offering a rich source of coding RNA data for model training. The diversity and volume of the mRNA sequences in RefSeq are instrumental in training a model that can generalize well across different species and biological contexts. Dataset statistics are shown in Table 2.

**Table 2:** Summary statistics of the pretraining datasets.

| Dataset | Number of Sequences | Total Nucleotides |
| --- | --- | --- |
| RNAcentral (ncRNA) | 36,000,000 | 26.42 billion |
| RefSeq (mRNA) | 532,852 | 2.22 billion |

## 3 Results

This section highlights the results from pretraining BiRNA-BERT and its performance on various downstream tasks. Section 3.1 covers the unsupervised clustering performance of BiRNA-BERT across different tasks including RNA secondary structural clustering, and RNA family clustering.

We performed a number of downstream tasks on varying sequence lengths, as shown in Figure 2. In Section 3.2, we demonstrate that BiRNA sets new state-of-the-art results on long sequence RNA-RNA interaction by leveraging dynamic tokenization which the previous language models handled with truncation.

**Figure 2:**
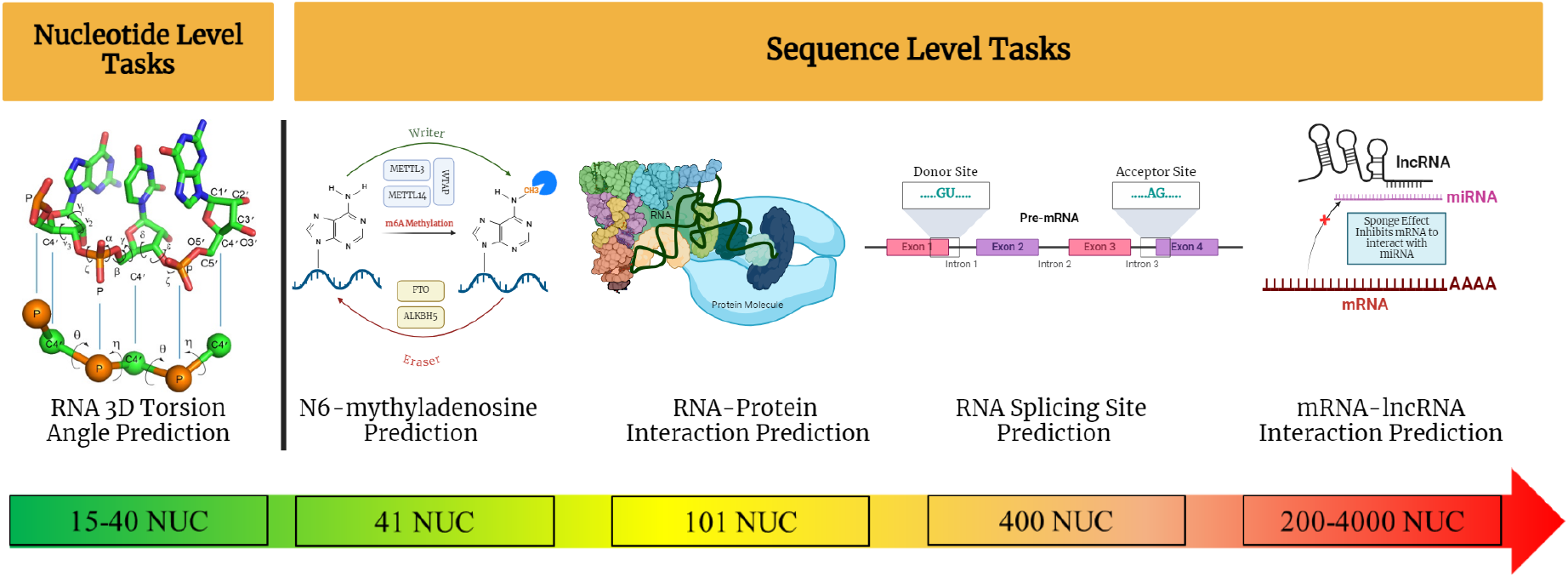
Downstream tasks for assessing the performance of various language models. The tasks are arranged from left to right according to increasing sequence length. We include one nucleotide-level task and four sequence-level tasks. The nucleotide-level task is RNA 3D torsion angle prediction. The sequence-level tasks are RNA N6-methyladenosine prediction, RNA-protein interaction prediction, RNA splicing site prediction, and mRNA-lncRNA interaction prediction.

We cover short sequence downstream tasks in Section 3.3 to establish that dual tokenization training has no drawbacks over conventional training. BiRNA-BERT even significantly surpasses similar-sized models such as RNA-FM and BERTRBP on short-sequence tasks. In Section 3.4, we demonstrate, on the RNA 3D torsion angle prediction task, that nucleotide embeddings from BiRNA have identical performance to a BERT trained solely on nucleotides. In section 3.5, we provide a rigorous mathematical analysis of the dual tokenization scheme from an information theoretic perspective which demonstrates the information loss from the BPE compression and how it compensates with the available GPU memory constraints. Moreover, we perform two different downstream tasks of RNA-Protein interaction and RNA N6-methyladenosine sites prediction to empirically show the validity of the information theoretic analysis.

We demonstrate, in section 3.6, the adaptability of our dual tokenization based approach on DNA sequences by training a smaller language model BiDNA-BERT for DNA sequences, which is 66X smaller architecture than the state of the art DNA language model DNABERT-2 [43]. Our results show that, despite being a small model, BiDNA-BERT provides comparable performance to DNABERT-2 on the human GLU benchmark.

### 3.1 Unsupervised Clustering Performance

To understand BiRNA-BERT’s capacity of unsupervised representation of the embedding space for RNA sequences from different structural families, we perform two different unsupervised clustering tasks. For the first task, we perform unsupervised clustering on 9 RNA structural families: 16s, 23s, 5s, RNaseP, grp1, srp, tRNA, telomerase, tmRNA. For the second task, we cluster the thirty RNA families from the rfam database. Here we compare unsupervised clustering performance between RNA-FM, RiNALMo and BiRNA-BERT.

After extracting the embeddings using BPE tokenization, we apply t-SNE for dimensionality reduction and visualize the 2D embedding space in Figure 3. As shown in the figure, BiRNA-BERT successfully clusters distinct RNA structural families as well as RNA Rfam families. To quantitatively assess the clustering performance of BiRNA-BERT in comparison to other methods on these tasks, we employ the silhouette coefficient as a measure of clustering quality. Silhouette coefficient is a metric used to evaluate the quality of clustering. It measures how similar a data point is to its own cluster compared to other clusters. The coefficient for a data point *i* is defined as:

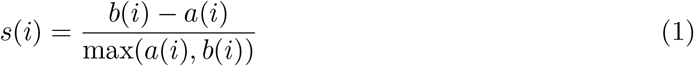

**Figure 3:**
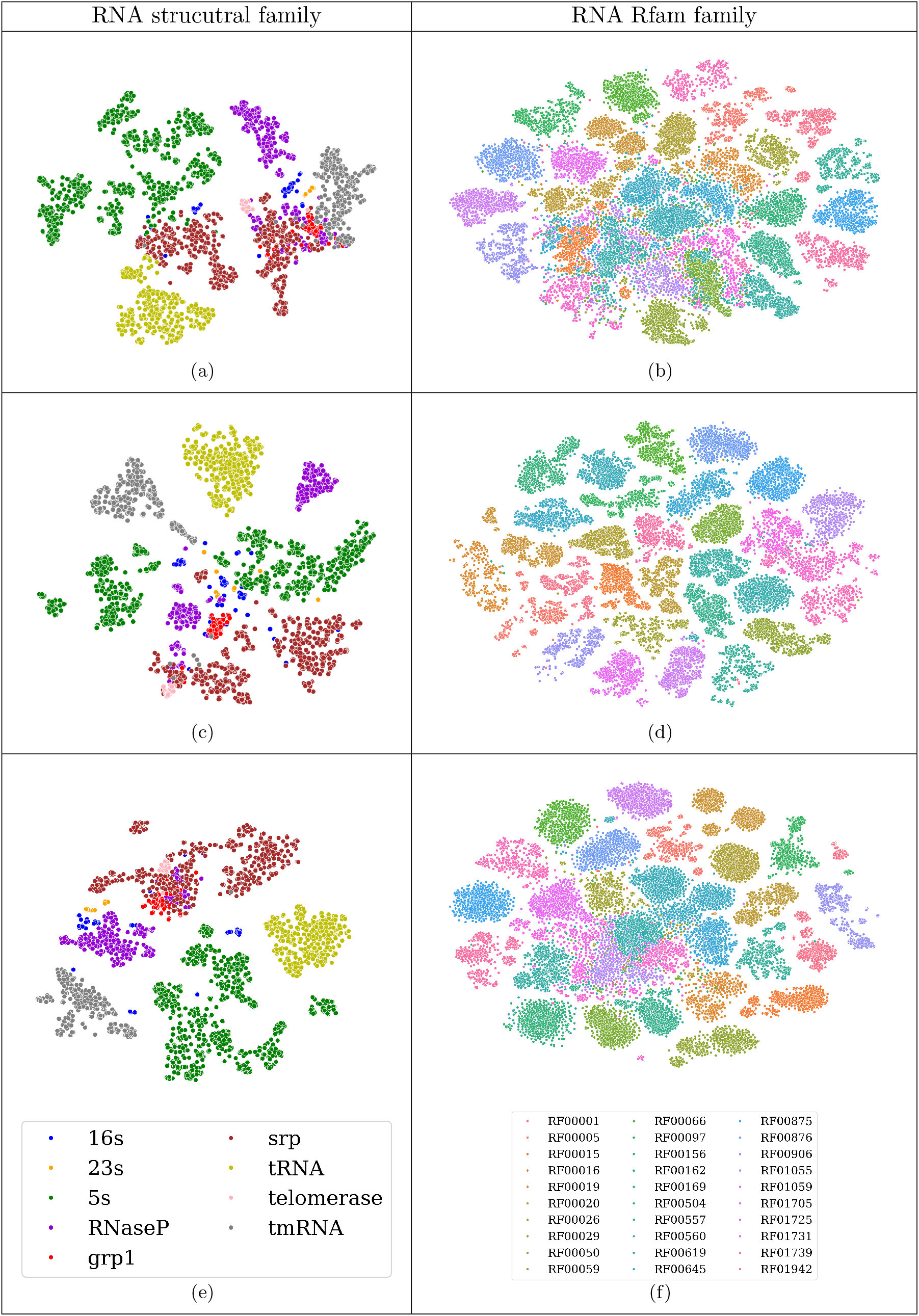
Comparison of unsupervised clustering performance on RNA structural families and RNA Rfam families. (a)-(b) RNA structural family and Rfam family clustering using RNA-FM. (c)-(d) Clustering using RiNALMo, and (e)-(f) clustering using BiRNA-BERT.

Here, *a*(*i*) represents the average distance between the data point *i* and all other points within the same cluster. In contrast, *b*(*i*) is the average distance between the data point *i* and all points in the nearest cluster to which *i* does not belong. The value of *s*(*i*) ranges from −1 to 1. A value of 1 indicates that the data point is well-matched to its own cluster and poorly matched to neighboring clusters. A value of 0 indicates that the data point is on or very close to the decision boundary between two neighboring clusters. Negative values suggest that the data point might have been assigned to the wrong cluster. A higher silhouette coefficient indicates well-defined clusters.

Table 3 shows that in the RNA structural family clustering task, BiRNA-BERT significantly outperformed both RNA-FM and RiNALMo. However, in the RNA Rfam clustering task, BiRNA-BERT significantly surpasses RNA-FM but slightly lags behind RiNALMo, with only a marginal difference in performance. Rfam dataset is one of the datasets in the RNAcentral database which has been used for pretraining the language models. Both RiNALMo and BiRNA-BERT use a masked language model approach, and thus a longer training duration is expected to enhance unsupervised clustering performance. Given that RiNALMo was trained for 14 days, while BiRNA-BERT was trained for only 2 days with a model six times smaller, it is understandable that BiRNA-BERT’s performance on the Rfam dataset would not surpass RiNALMo’s. However, the secondary structural family dataset was not explicitly included in the pretraining database. BiRNA-BERT significantly outperforms all larger language models on this task, demonstrating superior unsupervised language understanding compared to other models when applied to unseen datasets.

**Table 3:** Clustering performance (Silhouette Coefficient) of different models.

|  | RNA Structural Family Clustering | RFAM Family Clustering |
| --- | --- | --- |
| RNA-FM | 0.139 | 0.238 |
| RiNALMO | -0.001 | **0.377** |
| BiRNA-BERT | **0.156** | 0.351 |

### 3.2 Long-Sequence Task with Dynamic Tokenization: miRNA-lncRNA Interaction Prediction

Long non-coding RNA (lncRNA) related studies frequently involve much longer sequences which necessitated task-specific architectures such as PmliPred [17] and CORAIN [39]. We evaluate the interaction between lncRNA and micro RNA (miRNA) to verify the impact of sequence truncation during feature embedding by previous models and compare it with our approach, which avoids sequence cropping to fit computational memory. Instead, we dynamically compress sequence information using our adaptive tokenization scheme.

We use three benchmarking datasets for the RNA-RNA interaction prediction task compiled by [17] (dataset details shown in Table 4). For evaluation, one benchmarking dataset is used as the training set, and another dataset is used for validation following the strategy used in [39]. Thus, we have 6 train-test combinations and we report performance in all these combinations. The lengths of the sequences in the miRNA dataset are 10 to 50, whereas the lncRNA dataset ranges from 200 to 4000. The length distributions of sequences for the miRNA-lncRNA Interaction Prediction task are shown in Figure 4.

**Table 4:** Summary of the benchmark dataset for the miRNA-lncRNA interaction prediction task.

| Benchmarks of<br>RNA-RNA<br>interactions |  | No. of miRNAs | No. of lncRNAs | No. of<br>molecule pairs |
| --- | --- | --- | --- | --- |
| Arabidopsis thaliana<br>(ATH) | Interacting Pairs | 331 | 2014 | 2500 |
|  | Non-interacting<br>Pairs | 266 | 1964 | 2500 |
| Glycine max<br>(GMA) | Interacting Pairs | 401 | 1770 | 2500 |
|  | Non-interacting<br>Pairs | 542 | 171 | 2500 |
| Medicago truncatula<br>(MTR) | Interacting Pairs | 335 | 1986 | 2500 |
|  | Non-interacting<br>Pairs | 424 | 2442 | 2500 |

**Figure 4:**
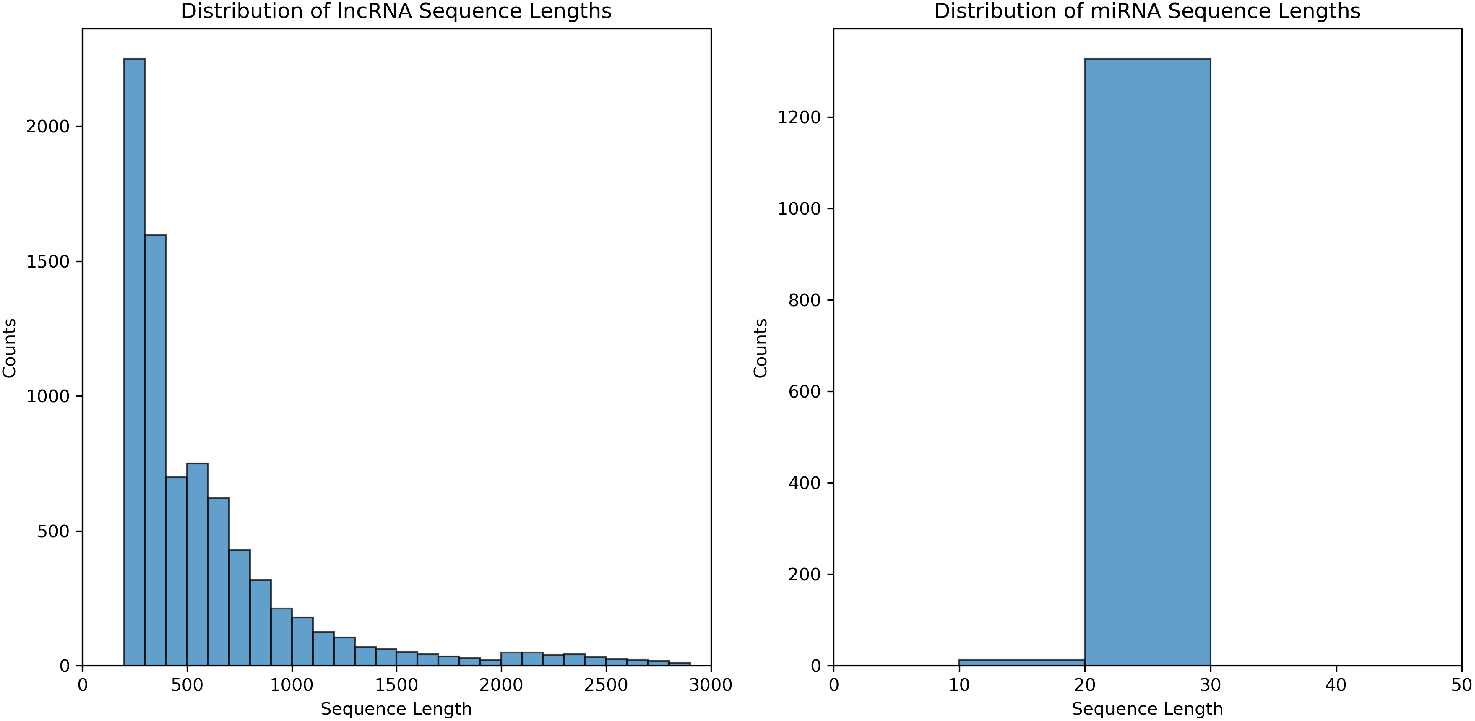
Distribution of sequence lengths in the lncRNA and miRNA datasets.

This is a binary classification task to determine whether a miRNA-lncRNA pair interacts or not. For fine-tuning, we follow the methodology of CORAIN [39] by freezing the backbone encoders, concatenating output features of miRNA and lncRNA, and using a simple convolutional prediction head. We test two strategies using BiRNA: BPE and NUC with truncation, named BiRNA-BPE and BiRNA-NUC. Both strategies use the same BiRNA models and only the input tokenization scheme differs. We always encode miRNA using NUC due to their short lengths. We use NUC with truncation for lncRNA for RNA-FM, RiNALMo, and BiRNA-NUC. We truncate all inputs to 1022 tokens due to the context window limitation of RNA-FM and RiNALMo even though BiRNA-NUC can process longer sequences due to ALiBi. We do not truncate input sequences to BiRNA-BPE since after BPE tokenization the maximum sequence length is 807, still lower than NUC.

The results in Table 5 offer several interesting insights:

**Table 5:** Accuracy of different models on miRNA-lncRNA dataset. CORAIN [39] is a task-specific CNN-autoencoder and the current state-of-the-art. We fine-tuned RNA-FM, RiNALMo, and both variants of BiRNA-BERT with optimal hyperparameters found using grid search.

| Model | Train-Test Dataset |  |  |  |  |  |
| --- | --- | --- | --- | --- | --- | --- |
|  | ATH-GMA | ATH-MTR | GMA-ATH | GMA-MTR | MTR-ATH | MTR-GMA |
| CORAIN | 69 | 74 | 67 | 93 | 58 | 84 |
| RNA-FM | 68.56 | 71.68 | 69.90 | 94.68 | 57.15 | 83.92 |
| RiNALMo | 72.14 | 75.38 | **70.93** | **95.08** | 65.55 | 85.82 |
| BiRNA-NUC | 72.56 | 77.90 | 66.01 | 94.94 | 61.05 | 84.18 |
| BiRNA-BPE | **75.52** | **79.56** | 70.55 | 93.84 | **67.75** | **88.54** |

1. RNA-FM performs significantly worse than the non-LM-based approach in 4 of 6 test datasets. However, RiNALMo outperforms the current state-of-the-art in all datasets despite sequence truncation. This is noteworthy, as RiNALMo is a six times larger language model than RNA-FM, significantly enhancing its expressive capability.
2. BiRNA-NUC outperforms RNA-FM in 5 out of 6 datasets and provides comparable performance to RiNALMo, despite being the same size as RNA-FM. BiRNA-BPE outperforms RiNALMo by a substantial margin, with improvements of 4.74%, 5.54%, 3.35%, and 3.16% on the ATH-GMA, ATH-MTR, GMA-MTR, and MTR-GMA datasets. It also offers comparable performance in the GMA-ATH and GMA-MTR datasets, within 0.51% and 1.2% margins.
3. An intuitive way to compare NUC and BPE tokenization is by considering information loss. NUC explicitly truncates sequences to 1022 nucleotides, losing all subsequent information. BPE, on the other hand, compresses the entire sequence (see Section 3.5). In miRNA-lncRNA interaction tasks, we demonstrate that compression (BPE) is preferable to explicit information loss (NUC), while also being more computationally efficient (807 tokens versus 1024 tokens).

BiRNA-BERT uses the information compression technique using BPE and encodes the long sequences with minimal information loss, thereby providing significant performance gain over the traditional models.

### 3.3 How Close in Short Sequences? : RNA Splicing Site Prediction

We consider the downstream tasks that can be performed within the computational limit of traditional BERT-based architecture as short-sequence tasks. We do not need to truncate the sequences in this case explicitly. We consider the task of binary classification of RNA splicing site prediction specifically for acceptor sites. RNA splicing is a crucial process in eukaryotic gene expression, where introns are removed from precursor messenger RNAs (pre-mRNAs), and exons are joined together to form mature mRNAs. This process is essential for generating functional mRNAs that can be translated into proteins. Identifying splice sites—the donor sites at the 5’ end of introns and the acceptor sites at the 3’ end—is vital for accurately predicting gene structure and location. For this task, we consider the dataset proposed by [25]. Particularly we use the gold standard dataset GS 1 which contains an equal number of positive and negative samples. The dataset consists of “confirmed” error-free splice-site sequences from a diverse set of 148 eukaryotic organisms, including humans. We have tested the performance of the trained model on three independent test datasets containing the samples from 3 different species of fish (*Danio rerio*), fruit fly (*Drosophila melanogaster*), and plant (*Arabidopsis thaliana*). Here the sequence length is 400 and the train and independent test tests have 20000 sequences each for training and testing respectively. We compare our model with RiNALMo, RNA-FM, and non-LM-based SOTA approaches in Table 6. BiRNA substantially outperforms Spliceator and the similar-sized RNA-FM (BERT) in all the datasets. Bi-RNA outperforms RiNALMo by 1.6% on the Fish dataset and is within 1.3% and 1.6% margins on Fly and Plant datasets.

**Table 6:**
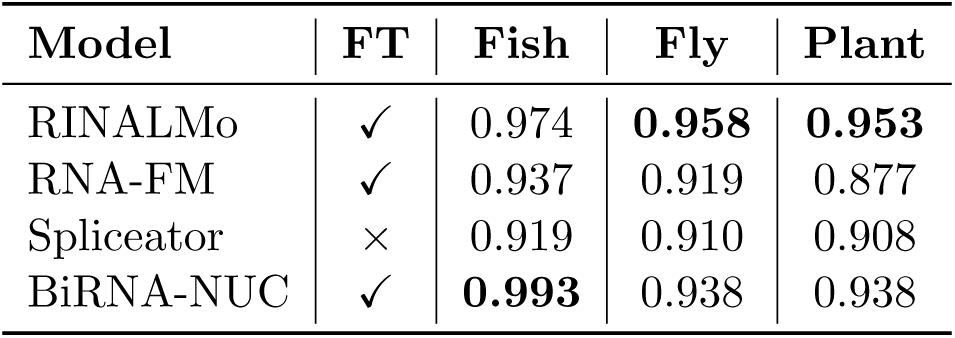
F1 score for RNA splicing site prediction on independent test sets. FT denotes whether the model uses finetuning or not.

### 3.4 Nucleotide-Level Task: 3D Torsion Angle Prediction

RNA 3D torsion angles are critical parameters that define the three-dimensional conformation of RNA molecules. In 3D torsion angle prediction, we need to predict 7 torsion angles for each nucleotide in a sequence: *α, β, γ, δ, ϵ, ζ*, and *χ*. These angles describe the rotations around the bonds that connect the nucleotides within an RNA strand, influencing its overall structure and stability. These are mathematically represented as the dihedral angles between four consecutive atoms in the RNA backbone. For example, the *α* angle is measured as the dihedral angle between O5’-P-O3’-C3’. The dataset for RNA torsion angle prediction is collected from https://sparks-lab.org/server/spot-rna-1d/. The training (TR), validation (VL), and three test sets (TS1, TS2, and TS3) have 286, 30, 63, 30, and 54 RNA chains, with average sequence lengths of 122, 15, 30, 14, and 24 respectively.

During finetuning, the prediction head outputs pairwise sine and cosine values for each angle, resulting in 14 output nodes. This is a regression task, and we minimize the mean squared error (MSE) between the predicted and actual sine-cosine pairs. We present the results in Table 7. BiRNA-BERT outperforms RNA-FM on all test datasets and matches closely with NUC-Only-BERT, our model trained using only NUC tokenization. This highlights that dual tokenization pretraining did not compromise the quality of nucleotide embeddings. The performance of both BiRNA-BERT and NUC-Only-BERT are comparable to RiNALMo 650M despite having only 117M parameters and being pretrained on 27× less compute. Thus, in addition to sequence-level tasks, the representations learned by BiRNA-BERT are effective for nucleotide-level tasks.

**Table 7:** Mean Absolute Error of RNA 3D torsion angle prediction. Here we report the mean squared error (MSE) between the predicted and actual RNA torsion angles. The “NUC-only” method refers to tokenization at the nucleotide level without any additional processing. We show the average error across different torsion angles for various methods. Model details are discussed in detail in Section 2.4. The best and second best results are shown in bold and italic, respectively.

| Data | Method | Avg Error | Alpha | Beta | Gamma | Delta | Epsilon | Zeta | Chi |
| --- | --- | --- | --- | --- | --- | --- | --- | --- | --- |
| VL | BPE-NUC | *28.085* | 29.132 | 24.018 | 25.692 | 21.345 | 25.718 | 35.918 | 34.771 |
|  | NUC-only | 28.398 | 29.583 | 23.910 | 26.195 | 21.779 | 26.193 | 35.948 | 35.177 |
|  | RNA-FM | 28.333 | 29.357 | 24.412 | 26.109 | 21.983 | 25.451 | 35.027 | 35.994 |
|  | RINALMo | **27.888** | 28.861 | 23.188 | 25.866 | 22.486 | 25.078 | 35.132 | 34.603 |
| TS1 | BPE-NUC | **28.181** | 30.223 | 21.856 | 19.553 | 29.193 | 34.232 | 26.764 | 35.449 |
|  | NUC-only | 28.760 | 31.002 | 22.887 | 19.793 | 29.415 | 34.607 | 27.980 | 35.637 |
|  | RNA-FM | 29.916 | 31.904 | 23.859 | 19.839 | 29.620 | 35.884 | 30.372 | 37.937 |
|  | RINALMo | *28.622* | 31.127 | 22.658 | 20.199 | 29.686 | 32.880 | 27.607 | 36.195 |
| TS2 | BPE-NUC | *26.704* | 24.728 | 17.363 | 23.836 | 19.104 | 31.875 | 36.048 | 33.973 |
|  | NUC-only | 27.252 | 25.602 | 18.408 | 24.406 | 19.748 | 32.139 | 36.474 | 33.990 |
|  | RNA-FM | 27.710 | 25.464 | 17.842 | 23.829 | 19.823 | 35.864 | 37.754 | 33.391 |
|  | RINALMo | **25.915** | 22.677 | 16.964 | 20.958 | 18.356 | 31.906 | 38.238 | 32.304 |
| TS3 | BPE-NUC | *31.979* | 34.728 | 17.363 | 23.836 | 19.104 | 31.875 | 36.048 | 33.973 |
|  | NUC-only | 32.174 | 34.856 | 19.606 | 30.324 | 37.531 | 36.589 | 35.374 | 30.938 |
|  | RNA-FM | 32.000 | 33.415 | 20.341 | 30.755 | 38.091 | 36.360 | 34.414 | 30.681 |
|  | RINALMo | **31.513** | 34.016 | 19.292 | 30.343 | 37.841 | 35.763 | 33.960 | 29.575 |

### 3.5 Information Theoretic Analysis of Nuelcotide vs BPE Tokenization

In this section, we compare the information content of a nucleotide token and a BPE token inspired by key empirical observations in the training data. Information-theoretic analysis of biological sequences is a well-studied field of research where the key challenges include determining the prior distribution of nucleotides or *k*-mers, the fact that only a fraction of possible biological sequences occurs in nature and the difficulty in comparing results from biological sequences with those from linguistics due to significant differences in morphology [1, 26, 37]. In addition to the information theoretic analysis, we further validate our analysis with two RNA downstream tasks.

#### Information content of a nucleotide token

We consider each nucleotide in a sequence as an independent variable that carries some amount of information. We wish to quantify the maximum amount of information for each new nucleotide in a sufficiently long sequence. We can derive the per-token upper bound of the Shannon entropy of a DNA/RNA sequence as follows.

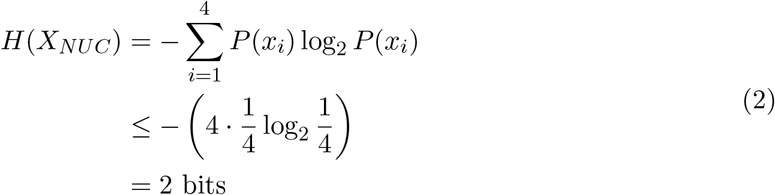

#### Information content of a BPE token

Similar to as nucleotide tokens, we consider each BPE token in a sequence as an independent variable that carries some amount of information. Let the size of the vocabulary be *N*. On our pretraining datasets, we observe that the frequency of the BPE tokens is exponentially distributed, and as a result, the probability of a token can modeled by an exponential function as follows.

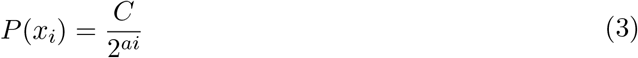

Since the index assigned to a token is arbitrary, tokens can be sorted by descending probability and reindexed without issue. Under this formulation:

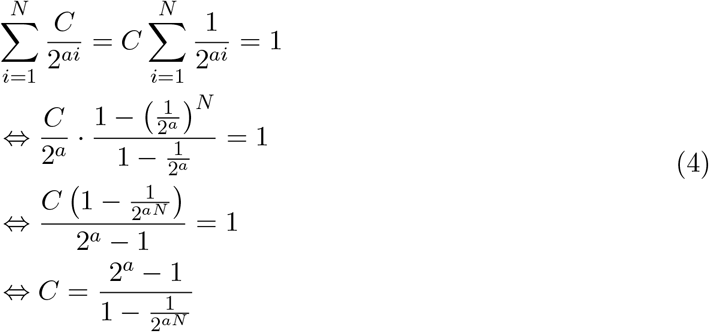

When the vocabulary size *N* is large, we can approximate *C* ≈ 2^*a*^ − 1 and *a* ≈ *log*_2_(*C* + 1). Now we can derive a general expression for the entropy of BPE tokens as,

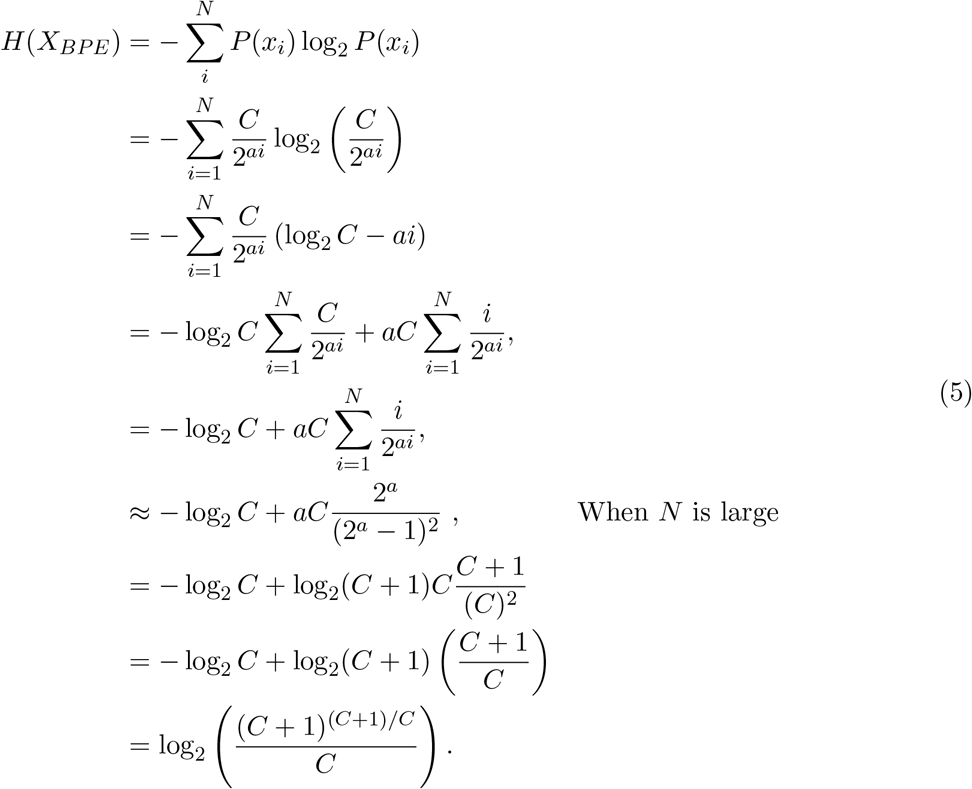

If the weighted average length of a BPE token is 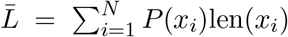, the average character-level entropy of BPE representation of a sequence will be 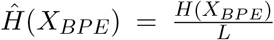. Since nucleotides are one character each, the per-character entropy is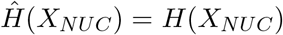.

The BPE tokenization will lead to less entropy if,

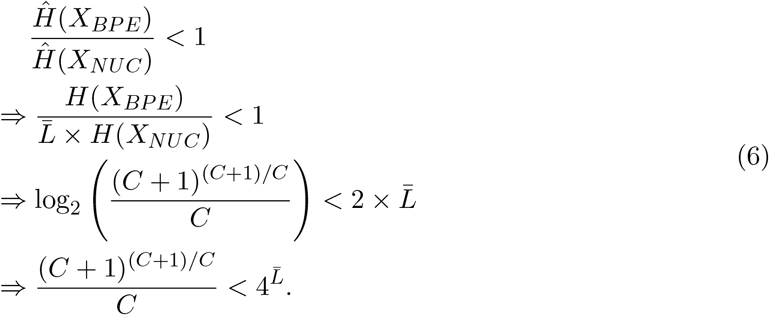

When *C* << 1, we can approximate (*C* + 1) ≈ 1. Then the inequality in Equation 6 can be further simplified as

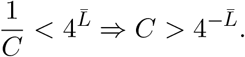

#### Empirical Entropy Ratio

On our pretraining data mixture, we determine that *P* (*A*) ≈ 0.2726, *P* (*A*) ≈ 0.2144, *P* (*A*) ≈ 0.26642, *P* (*A*) ≈ 0.2465, and averge BPE token length 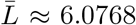. This yields the empirical entropy of nucleotide tokens *H*_*e*_(*X*_*NUC*_) ≈ 1.9939 bits. As shown in Figure 5, the empirical value of C is 0.005086 when determined on 33 million sequences of our pretraining dataset.

**Figure 5:**
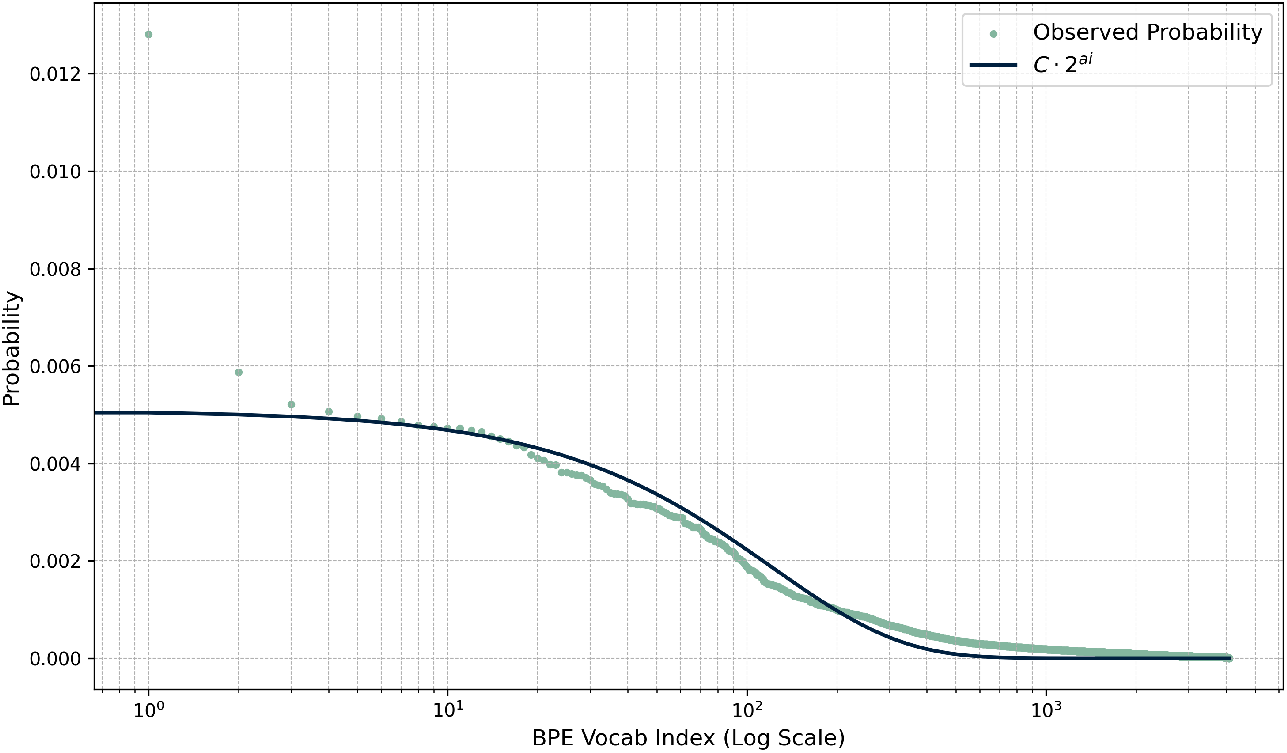
Fitting the exponential function *C ·* 2^−*ai*^ to the empirically observed BPE token probabilities on pretraining datasets. Since the index assigned to a token is arbitrary, we sort the tokens in an descending order of the probability/frequency and re-index tokens to represent the probability as a function of the index. We determined best-fit when *C* ≈ 0.005086 and *a* ≈ 0.011909.

Plugging in this value in Eqn. 5 yields *H*_*e*_(*X*_*BPE*_) ≈ 9.1044 bits. Therefore, the empirical per-character entropy ratio is

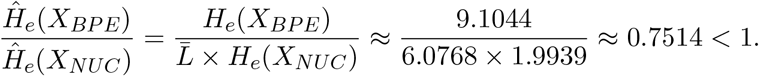

The empirical per-character entropy ratio of 0.7514 indicates that the BPE tokenization technique effectively compresses the input sequence. Although compressed information is likely more difficult for language models to process, it is well-compensated by the ability to process sequences up to 6 times longer than the original input with the same GPU memory constraints. This also partially explains why we observed BPE underperforming their NUC counterparts on short-sequence downstream tasks from an Information-theoretic perspective.

Therefore, BPE tokenization is essentially a trade-off between information compression and computational efficiency, which BiRNA-BERT can dynamically adjust depending on the hardware constraints and sequence length.

Here, we assume tokens are independent and identically distributed random variables (i.i.d) to approximate the information content of NUC and BPE sequences. In reality, the information content of non-i.i.d sequences is much lower than Shannon Entropy due to the correlation between nearby symbols. Language entropy [29] and Kolmogorov Complexity [18] take symbol correlation and order into account but are generally intractable.

#### Advantages of NUC over BPE for Short Sequences

To further validate our information theoretic analysis, we present a comparative performance analysis between BPE and NUC tokenization on short sequences across two downstream binary classification tasks: i) *RNA-Protein interaction prediction* and ii) *RNA N6-methyladenosine site prediction*. RNA-Protein interaction focuses on finding the binding sites and interactions between RNA molecules and proteins to understand post-transcriptional regulation. Benchmark dataset for RNA-protein interaction prediction is collected from RBPsuit available at http://www.csbio.sjtu.edu.cn/bioinf/RBPsuite/. This database contains datasets for 154 different proteins and a collection of interacting and non-interacting RNA sequences for each protein. We consider a subset of 5 datasets for our evaluation (AARS, AATF, AKAP1, AGGF1, ABCF1). The length of RNA sequences used for this task is 101 across all the datasets.

Task 2 (RNA N6-methyladenosine site prediction) focuses on predicting N6-methyladenosine (m6A) sites. N6-methyladenosine is a common and critical modification in eukaryotic mRNA, affecting various aspects of RNA metabolism. This includes stability, splicing, and translation. The prediction and detection of m6A sites are essential for understanding how this modification influences gene expression and cellular processes. In our work, we utilized datasets from human, rat, and mouse tissues, specifically focusing on brain, kidney, and liver samples for each species. Data sets were derived from the iRNA-m6A study available at http://www.biolscience.cn/Deepm6A-MT/data/, employing an antibody-independent m6A-REF-seq protocol, which is both high-throughput and accurate for m6A detection. Positive samples were selected based on the presence of m6A at the center of 41 continuous nucleotide residues, while negative samples were randomly selected from the same tissues but without m6A sites.

In Table 8, we present results on RNA-Protein interaction prediction for BPE and NUC tokenization under two different configurations of BiRNA-BERT which we described in Section 2.4. NUC consistently outperforms BPE on all five concerned datasets by 3.56% (AARS), 2.91% (AATF), 4.15% (ABCF1), 2.35% (AGGF1), and 3.50% (AKAP1). BiRNA outperforms the current SOTA (BERTRBP [40]) by 3.49% (AARS), 2.89% (AATF), 7.43% (ABCF1), 3.84% (AGGF1), and 4.10% (AKAP1). BERTRBP is explicitly trained for predicting RNA-Protein binding prediction task finetuned on DNABERT [15]. Results for RNA N6-methyladenosine site prediction are presented in Table 9. Similar to the RNA-Protein interaction task, NUC versions perform consistently better than BPE versions. The average improvement of NUC over BPE for human, mouse and rat are 3.88%, 2.24%, and 1.68% across three different tissues: brain, kidney, and liver. Moreover, BiRNA outperforms the current SOTA (Deepm6A-MT [14]) by 0.33% (Human), 0.45% (Mouse), and 0.06% (Rat).

**Table 8:** F1 score of different methods for RNA-Protein interaction prediction.

| Methods | Model | Datasets |  |  |  |  |
| --- | --- | --- | --- | --- | --- | --- |
|  |  | AARS | AATF | ABCF1 | AGGF1 | AKAP1 |
| BPE | BPE-Only | 0.6792 | 0.7014 | 0.7266 | 0.6946 | 0.6778 |
|  | BiRNA-BPE | 0.6769 | 0.6931 | 0.7296 | 0.6960 | 0.6835 |
| NUC | NUC-only | 0.6964 | 0.7217 | **0.7599** | 0.7101 | 0.7030 |
|  | BiRNA-NUC | **0.7034** | **0.7218** | 0.7528 | **0.7124** | **0.7074** |
|  | BERT-RBP | 0.6797 | 0.7016 | 0.7066 | 0.6861 | 0.6795 |

**Table 9:** Accuracy for RNA N6-methyladenosine sites prediction on various tissues across different organisms.

| Species | Human |  |  | Mouse |  |  | Rat |  |  |
| --- | --- | --- | --- | --- | --- | --- | --- | --- | --- |
| Tissue | Brain | Kidney | Liver | Brain | Kidney | Liver | Brain | Kidney | Liver |
| **BPE-Only** | 0.710 | 0.778 | 0.786 | 0.773 | 0.792 | 0.708 | 0.764 | 0.817 | 0.815 |
| **BiRNA-BPE** | 0.714 | 0.786 | 0.795 | 0.779 | 0.794 | 0.709 | 0.769 | 0.817 | 0.818 |
| **NUC-Only** | **0.753** | 0.807 | **0.823** | **0.802** | 0.816 | 0.745 | **0.788** | 0.831 | 0.824 |
| **BiRNA-NUC** | 0.747 | **0.811** | 0.816 | 0.801 | **0.820** | 0.747 | 0.785 | 0.831 | **0.825** |
| **Deepm6A-MT** | 0.751 | 0.809 | 0.815 | 0.799 | 0.816 | 0.733 | 0.785 | **0.840** | 0.818 |

### 3.6 Beyond RNA Sequences: BiDNA-BERT

We have extended the concept of dual tokenization with AliBi encoding to DNA sequences (which we call BiDNA-BERT) to evaluate the adaptability of our proposed scheme to other biological language models beyond RNA. To assess performance, we compared BiDNA with DNABERT-2 [43], the current state-of-the-art in DNA language modeling. While BiDNA-BERT was trained on the human genome using 3 million DNA sequences, DNABERT-2 was trained on a multi-species genomic dataset covering genomes from 135 species, nearly 12 times the volume of the human genome dataset. We evaluated all three human-genome-related downstream tasks from the GUE benchmark [43]. The tasks are as follows:

1. **Promoter Detection (Human):** Sequences were extracted from −249 to +50 bp around the TSS, categorized into TATA and non-TATA classes based on the presence of a TATA box. The TATA box is a specific DNA sequence found in the promoter region of many genes, typically located about 25-35 base pairs upstream of the transcription start site (TSS). It has a consensus sequence of “TATAAA” and is a crucial element for initiating the transcription process in eukaryotic cells.
2. **Core Promoter Detection (Human):** A narrower window of −34 to +35 bp around the TSS was used to target the core promoter region. This shorter context emphasizes the challenge of precise core promoter identification.
3. **Transcription Factor (TF) Binding Site Prediction (Human):** Data from 690 ENCODE ChIP-seq experiments covering 161 TF binding profiles were used. A 101-bp region around peak centers was selected, with non-overlapping sequences curated to balance task difficulty.

As shown in Table 10, BiDNA with NUC tokenization performs comparably to DNABERT-2 in the promoter site detection task, with only a 0.8% difference in performance. In the core promoter site detection task, BiDNA-NUC outperforms DNABERT-2, achieving a 4.4% higher MCC. For the transcription factor binding site task (Table 11), BiDNA demonstrates competitive performance, with a margin of just 1.2%. Notably, BiDNA achieves these results while using 66 times less computational resources than DNABERT-2.

**Table 10:** Comparison of BiDNA-BERT with DNABERT-2 for the promoter site and core promoter site prediction tasks. We show Mathews Correlation Coefficient (MCC) scores for various methods.

| Task | Promoter site prediction |  |  | Core promoter site detection |  |  |
| --- | --- | --- | --- | --- | --- | --- |
| Dataset | All | No TATA | TATA | All | No TATA | TATA |
| BPE-Only | 0.916 | 0.954 | 0.799 | 0.806 | 0.824 | 0.765 |
| BiDNA-BPE | 0.918 | 0.954 | 0.789 | 0.812 | 0.828 | 0.758 |
| NUC-Only | 0.927 | 0.963 | 0.807 | **0.832** | 0.836 | **0.876** |
| BiDNA-NUC | 0.936 | 0.966 | 0.817 | **0.832** | 0.839 | 0.844 |
| DNABERT-2 | **0.941** | **0.971** | **0.830** | 0.831 | **0.849** | 0.814 |

**Table 11:**
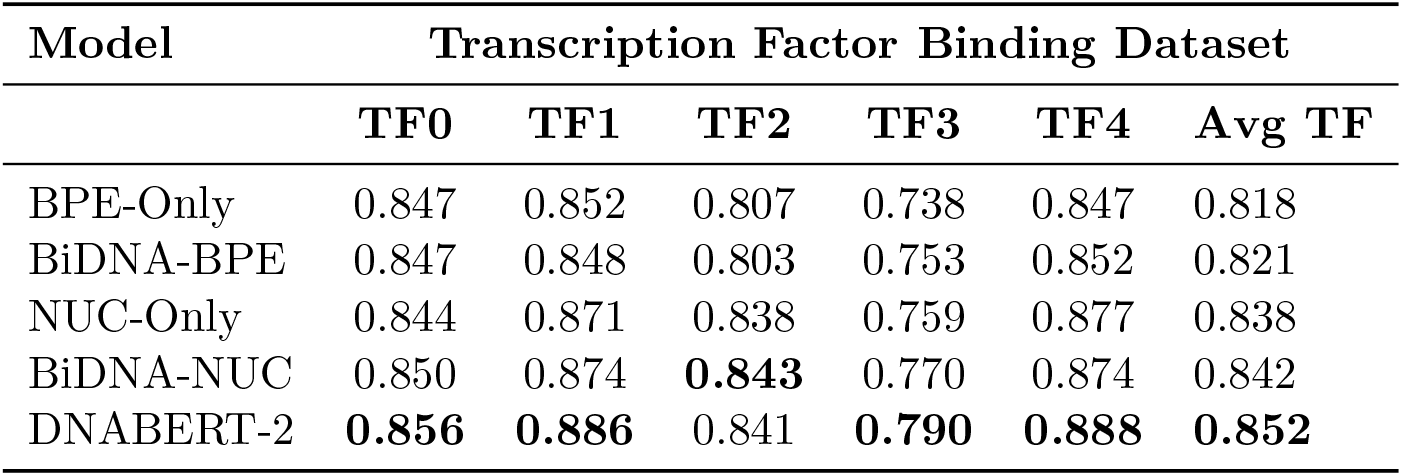
Comparison (in terms of MCC scores) of BiDNA-BERT with DNABERT-2 for the transcription factor binding prediction task across different datasets.

| Model | Transcription Factor Binding Dataset |  |  |  |  |  |
| --- | --- | --- | --- | --- | --- | --- |
|  | TF0 | TF1 | TF2 | TF3 | TF4 | Avg TF |
| BPE-Only | 0.847 | 0.852 | 0.807 | 0.738 | 0.847 | 0.818 |
| BiDNA-BPE | 0.847 | 0.848 | 0.803 | 0.753 | 0.852 | 0.821 |
| NUC-Only | 0.844 | 0.871 | 0.838 | 0.759 | 0.877 | 0.838 |
| BiDNA-NUC | 0.850 | 0.874 | **0.843** | 0.770 | 0.874 | 0.842 |
| DNABERT-2 | **0.856** | **0.886** | 0.841 | **0.790** | **0.888** | **0.852** |

## 4 Conclusions

The application of NLP techniques to biological research has seen renewed interest following the success of AlphaFold [16] in predicting protein structures. While the majority of research efforts have focused on protein sequences with considerable success, recent interest has begun to shift towards modeling DNA and RNA sequences. Bioinformatics researchers aim to develop foundation models that can be fine-tuned for a range of downstream tasks where labeled data is limited. The existing foundation models in DNA/RNA sequences cannot handle long sequences. However, we have a number of tasks with long sequences, for instance, the mRNA sequence database on NCBI [27] has an average length of 4,175 nucleotides. Additionally, tasks like secondary structure prediction [35] and torsion angle prediction [32] often require generating features at the nucleotide level. Hence, it is crucial to manage longer sequences without compromising nucleotide-level analysis, and our proposed dual tokenization balances this trade-off effectively.

This study first empirically demonstrates that two popular tokenization approaches, NUC and BPE, each have their own advantages in RNA sequencing tasks: BPE, along with ALiBi positional encoding, enables transformer encoder models to process long biological sequences, while NUC enhances predictions for high-granularity tasks, yielding performance gains over shorter sequences. Given these observations, we propose a dual tokenization approach and show that pretraining on both NUC and BPE tokenizations of a sequence allows a single model to support both with no downsides compared to training on either alone. We release a new RNA foundational model, BiRNA-BERT, trained over our proposed tokenization approach, which achieves state-of-the-art results in long-sequence tasks and outperforms similar-sized models in short-sequence and nucleotide-level tasks. We also validate our methodology on DNA data and provide an information-theoretic analysis comparing NUC and BPE tokenization. Our information theoretic analyses show that, when the sequence lengths are small enough to be handled by the available GPU memory, nucleotide tokenization consistently outperforms the BPE compression scheme – supporting our proposal of using the adaptive tokenization scheme (Nucleotide tokenization for short sequences and shifting to BPE compression for much longer sequences).

Our current approach reflects the simplest design of combining NUC and BPE. This naturally led to BPE tokens being underrepresented in the pretraining data: one BPE token corresponds to 6 nucleotides on average. The quality of embeddings generated with BPE tokens might improve by oversampling BPE tokens. BiRNA-BERT is computationally magnitude times (27X) smaller architecture than the existing largest RNA language model RiNALMo [21]. Moreover, it was trained for only 1 epoch, which is significantly shorter than other biological sequence models. Despite its constrained architecture, BiRNA-BERT achieves state-of-the-art results in multiple downstream tasks. It significantly outperforms other foundation models in long sequence downstream tasks, which have remained among the most challenging in biological language modeling to date. Furthermore, in unsupervised clustering tasks, BiRNA-BERT demonstrates superior performance compared to current state-of-the-art models, highlighting its enhanced ability to understand the complex language of RNA genomics. We invite the research community to improve upon our findings with larger foundation models. Our proposed approach generalizes across various biomolecular sequences, including DNA, RNA, and proteins. We extended this framework to create BiDNA-BERT, a small model trained on only 3 million DNA sequences, which achieved competitive results when compared to significantly larger DNA foundation models like DNABERT-2. The performance of BiDNA-BERT can be further improved in a straightforward way by increasing the language model size as as downstream performance is directly aligned with the model size of the BERT based language models [31]. Our language model can be similarly adapted for protein sequences. Thus, future studies need to leverage our proposed model to build foundation models for various biological sequences, thereby offering an effective solution for handling arbitrarily long sequences without sacrificing nucleotide-level granularity. We opted for ALiBi over competition methods such as RoPE [34] since ALiBi supports dynamically increasing the context window without retraining. Although BPE is prevalent in NLP, it has its drawbacks [2]. Exploring dual tokenization schemes with varied positional encodings and algorithms holds promise for future research.

## 5 Data and Code Availability

The datasets underlying this article were derived from sources in the public domain. The code and data are available at: https://github.com/buetnlpbio/BiRNA-BERT.

## Footnotes

1 The code and model weights are available at https://github.com/buetnlpbio/BiRNA-BERT

2 https://ftp.ncbi.nlm.nih.gov/refseq/release/

